# Intra- and Inter-individual genetic variation in human ribosomal RNAs

**DOI:** 10.1101/118760

**Authors:** Artem Babaian

## Abstract

The ribosome is an ancient RNA-protein complex essential for translating DNA to protein. At its core are the ribosomal RNAs (rRNAs), the most abundant RNA in the cell. To support high levels of transcription, repetitive arrays of ribosomal DNA (rDNA) are necessary and the long-standing hypothesis is that they undergo sequence homogenization towards rDNA uniformity.

Here I present evidence of the rich genetic diversity in human rDNA, both within and between individuals. Using state-of-the-art genome sequencing data revealed an average of 192.7 intra-individual variants, including some deeply penetrating the rDNA copies, such as the bi-allelically expressed 28S.59A>G. From 104 diverse genomes, 947 high-confidence variants were identified and unmask a hidden genetic diversity of humans.

These findings support the emerging concept that ribosomes are heterogeneous within cells and extends the heterogeneity into the realm of population genetics. Fundamentally, do our ribosomal variants determine how our cells interpret the genome?

## Introduction

The ribosome is an RNA-protein complex essential to all life. This ancient complex translates genetic information into protein. In humans, a mature ribosome contains four structural and catalytic RNA molecules: 18S in the small subunit and 5S, 5.8S and 28S in the large subunit.

Ribosomes are highly abundant in cells with ribosomal RNA (rRNA) making up >80% of the RNA in a human cell.(1) To be transcribed to such a high level, the DNA encoding for rRNA (rDNA) is present at ~600 copies per diploid genome. The *RNA45S* gene encodes the 45S pre-rRNA, which is processed and modified into 18S, 5.8S and 28S. The 13.5 kb *RNA45S* gene sits within a 43 kb rDNA repeat, which in turn are arranged in tandem arrays making up the acrocentric arms of chromosomes 13, 14, 15, 21 and 22. The 5S rRNA gene, RNA5S, is found on a tandem-repeated locus at chromosome 1p42.13.(2)

In the initial human genome sequencing project rDNA-containing BACmids were systematically depleted since sequencing was expensive and these repetitive sequences are thought to ’homogenize’ into a uniform sequence.(3) Yet notably, in the original research yielding the reference human rDNA, Kuo and colleagues repeatedly cloned and sequenced variants both within and between individuals.(4,5) The variation in these clones localized to ’variable loop regions’ and at the time it would have been challenging to distinguish these variants from pseudogenized rDNA. With recent improvements in sequencing technology, rDNA variation has been increasingly characterized in a diverse collection of species(6-10),so rRNA may not be as homogeneous as previously thought.

Originally I hypothesized that a transposable element could exist within human rDNA (similar to the *Drosophila* R1 and R2 elements(11)) yet would be undetected by standard variant analysis of the human genome. I interrogated human rDNA next-generation sequencing data and ’re-discovered’ massive intra- and inter-individual sequence heterogeneity in human rDNA, including expressed variants of 18S and 28S.

## Results

### Inter-individual variation in rDNA

To measure the intra-individual variation in rDNA, high coverage PCR-free whole genome sequencing data(12) (DNAseq) from the Utah CEPH-1463 trio (daughter, NA12878; father, NA12891; mother, NA12892) was aligned to a reference human rDNA sequence, *hgr1* (see materials and methods and Figure 1A).

**Figure 1:**
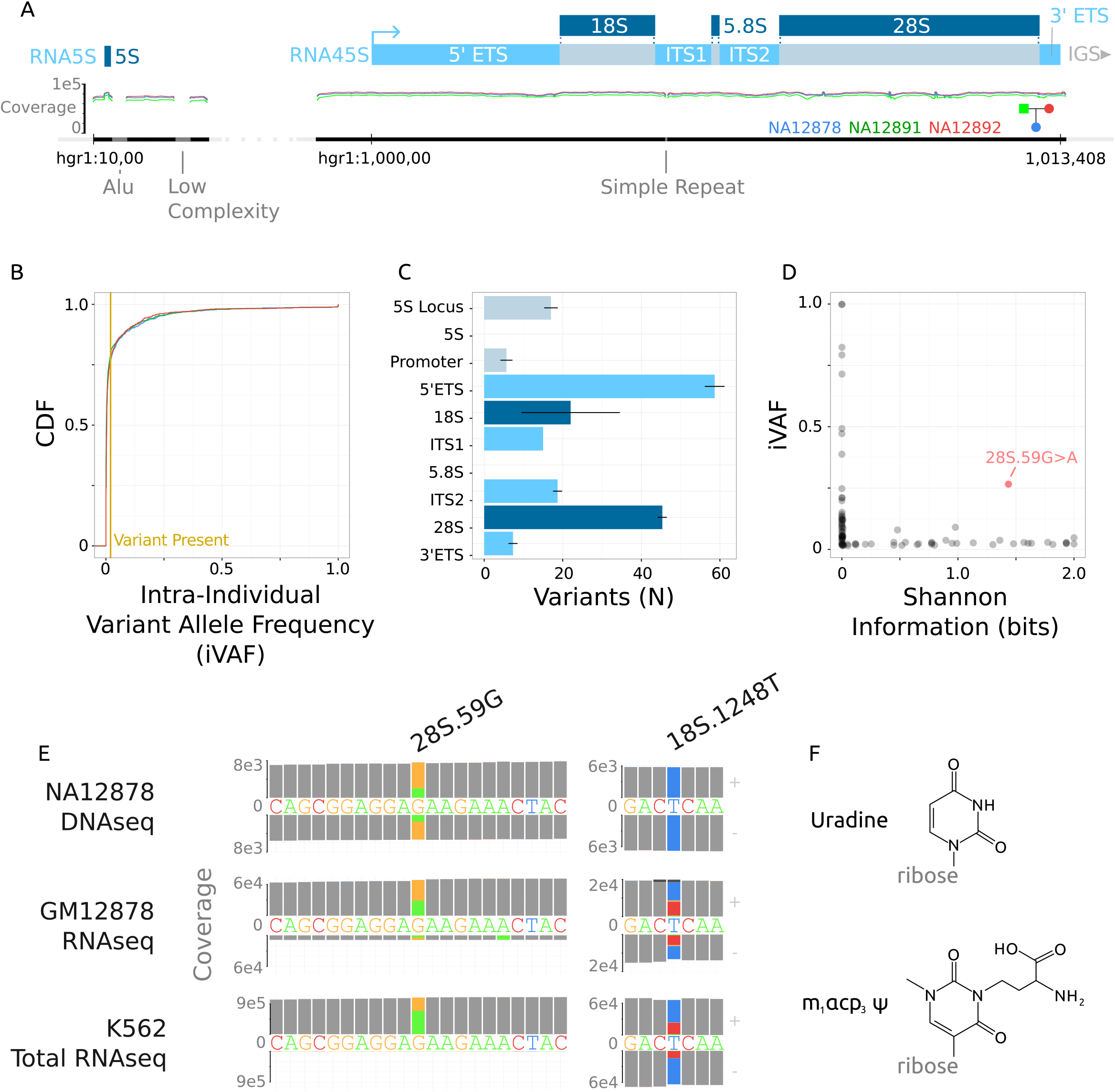
Intra-individual variation at the rDNA locus. A) Map of the *hgr1* reference sequence used in this study. It includes a single repeat of the entire *RNA5S* locus and *RNA45S* with adjacent sequences. In each locus a simple-repeats and a AluY were N-masked. *RNA45S* can be subdivided into the 5’ and 3’ external transcribed spacer (ETS), internal transcribed spacer (ITS) 1 and 2, and the mature rRNAs. Downstream of each *RNA45S* is a 29.6kb intergenic spacer (IGS) which was not included in this study. B) Log-scaled coverage across *hgr1* in the CEPH-1463 trio. C) The cumulative distribution of intra-individual variant allele frequency of the 926 variant positions (see materials and methods) for each member of the trio. The yellow vertical line at iVAF = 0.02 is the threshold at which a variant was called as ’present’ in an individual. D) The average distribution of variants (iVAF > 0.02) in each region of *hgr1.* E) Evolutionary conservation (in Shannon Information bits(26)) of the mature rRNA variants relating to the intra-individual variant allele frequency. F) Detailed coverage of NA12878 DNA- and poly-A selected RNA-seq, as well as K562 total RNA-seq at 28S.59G and 18S.1248T. The read coverage matching the reference sequence is colored in gray and mis-matching sequences are colored by the alternative base. E) The structure of i) uradine and ii) the hyper-modified nucleotide 1-methyl-3-(α-amino-α-carboxyl-propyl) Psuedouridine.

*hgr1* was covered to an average depth of 8853x (Figure 1B). There are an average of 192.7 (+-14.3) variants per individual at an intra-individual variant allele frequency (iVAF) of greater than 2%. With a 10% iVAF cut-off, there are 85.3 (+- 4.2) variants per individual (Figure 1C). The distribution of the variants is uneven over the different regions of rDNA (Figure 1D). Excluding 5S and 5.8S which were invariable, the density of variation was lower in 18S and 28S at 9.7 variants per kilobase (kb) compared to the other transcribed regions at 15.8 variants per kb, a trend consistent with rDNA variation in other species.(8,10)

As expected, the evolutionary conservation of the mature rRNA variants were at evolutionarily variable regions, such as the expansion loops, or were rare within individuals. The outlier to this trend is the 28S.59G>A variation, which is both evolutionarily conserved and highly variable within individuals. (Figure 1D).

To test if the variants in the mature rRNA are expressed, poly-A selected RNA sequencing (RNAseq) from the ymphoblastoid cell line GM12878 (derived from NA12878) was analyzed.(13) Variants at positions of moderate GC-content could be analyzed such as 28S.59G>A which shows bi-allelic expression. To interrogate a purer rRNA library, K562 total RNAseq (not poly-A selected and not rRNA-depleted) was analyzed with 70.3% of all reads mapping to rDNA, and here too, 28S.59G>A shows bi-allelic expression (Figure 1E).

While inspecting RNAseq libraries, one variant was recurrently observed at the highly conserved 18S.1248T, yet was absent from DNAseq. In the reference RNA, this base is a uradine, but in the mature 18S rRNA, this base is hyper-modified to 1-methyl-3-(α-amino-α-carboxyl-propyl) Pseudouridine (Figure 1E) for its function in the catalyitic core of the ribosome’s P-site. This hyper-modification hinders reverse transcriptase function(14), thus the "variation" is likely an error profile specific to this modified nucleotide. Besides being interesting, these findings indicate that most standard poly-A selected RNAseq libraries have ‘contaminating’ levels of rRNA which can be used to measure variant rRNA expression in future studies.

### Inter-individual variation in rDNA

To measure if there is rDNA variation between individuals, an additional 104 genomes (four genomes from each of 26 populations)(15) were aligned to *hgr1*.

Combined variant discovery (see materials and methods) from all 107 genomes yielded 947 variants at 926 distinct positions passing quality control (p < 0.001) (Figure 2A, Supplementary File 1). 78% of variants are single base substitutions favoring GC base changes and the remaining are near-equally divided between short insertions and deletions (Figure 2B).

**Figure 2:**
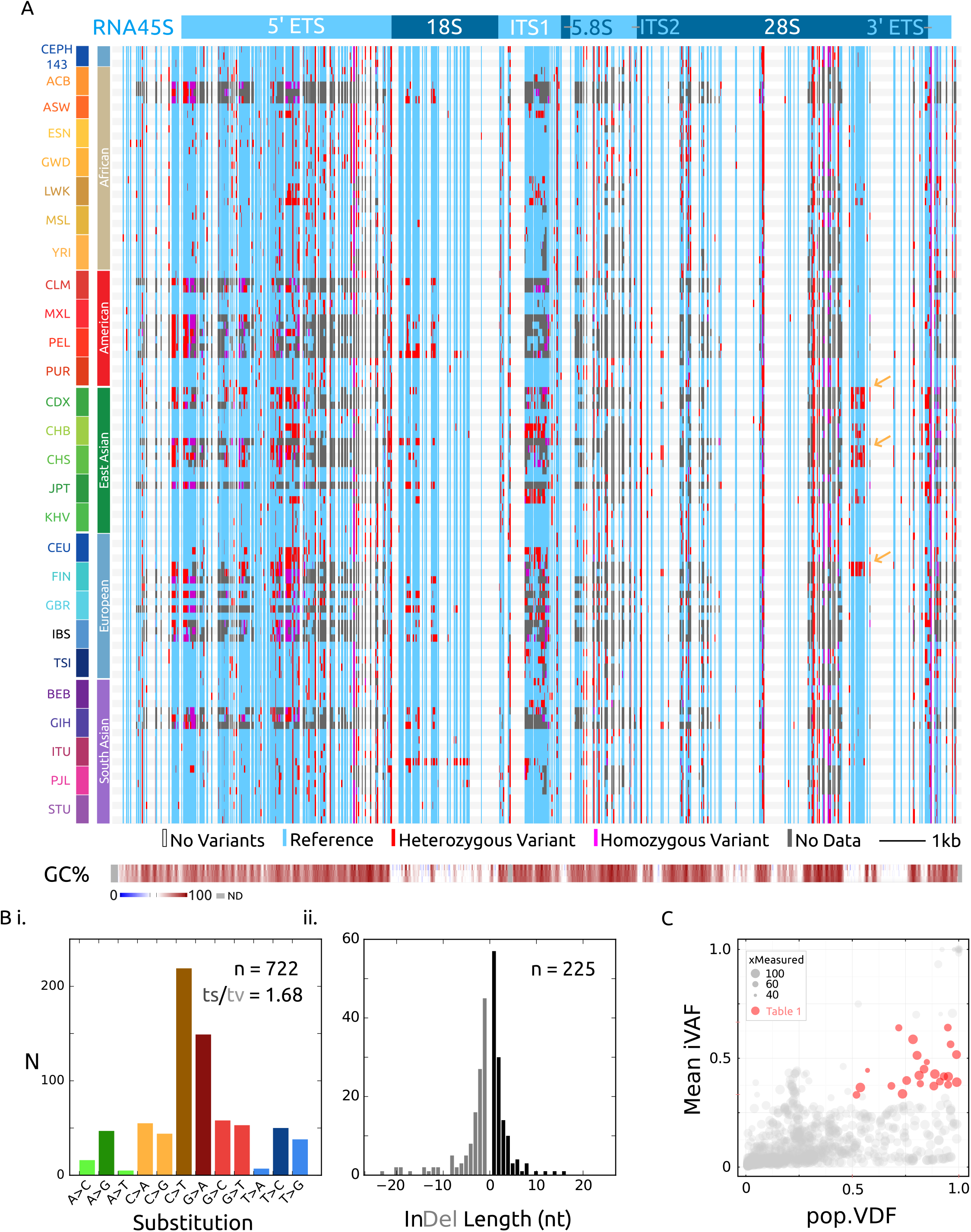
Inter-individual variation at *RNA45S*. A) Visualization of the high-quality variants from 107 individuals from 26 world populations. Vertical blue lines are sequences in which a variant genotype was called in at least one genome, with heterozygous and homozygous genotyping in red and magenta respectively. Invariable positions are clear columns across the whole graph. Particular libraries in which there was no quality coverage at a variant is gray. Sequence GC% is shown at the bottom in 30, 50 and 70 bp sliding windows. F) Transition vs. transversion nucleotide substitutions in the 722 single-nucleotide variants. E) Size distribution of the 225 indel variants. D) Inter-individual, or population variant-detection frequency (pVDF) vs. Intraindividual variant allele frequency (iVAF) scaled by the number of distinct genomes in which the variant allele was measured (to account for regions of uneven read coverage in the data). Red highlighted variants have an average intra-individual allele frequency between 0.33 - 0.66 and are found in at least half of the genomes interrogated (see Table 1).

The density of variation across *RNA45S* largely matches the CEPH-1463 trio distribution but offers finer resolution of regions absent of variation, notably 5S and 5.8S, and regions of 18S and 28S. It is important to note that regions of extreme GC-content had lower sequencing coverage across the DNAseq libraries, and more ’no data’ points then moderate-GC regions, something which can be addressed with deeper sequencing libraries in the future.

Reasoning that biologically relevant variants, with respect to cell- or tissue-heterogeneous ribosomes, would be maintained in humans, I quantitatively assessed the distribution of the variants in the population (Figure 2C). There is a cluster of 23 variants which are both, prevalent in the population, and abundant within individuals (population variant-detection frequency (pVDF) > 0.5, iVAF 0.33 - 0.66). Alternatively these variants could have had a higher iVAF in human ancestors, and drift could explain their prevalence and abundance as well. To truly measure balancing selection in rDNA, a more complete understanding of its heredity is necessary, but these 23 candidate variants warrant priority biochemical and molecular analysis (Table 1).

**Table 1:** Prevalent and abundant rDNA variants

| Coordinates (hgr1:) | Region | Reference Allele | Variant Alleles | iVAF | sd(iVAF) | pVDF | Variant Calls (N) |
| --- | --- | --- | --- | --- | --- | --- | --- |
| 1000777 | 5' ETS | CGG | C, CGGG, CGGGG | 0.52 | 0.29 | 0.99 | 94 |
| 1001683 | 5' ETS | G | GT | 0.37 | 0.19 | 0.88 | 83 |
| 1001756 | 5' ETS | C | T | 0.45 | 0.17 | 0.84 | 77 |
| 1001944 | 5' ETS | G | C, CGGG | 0.42 | 0.26 | 0.95 | 97 |
| 1002492 | 5' ETS | TGCG | T | 0.64 | 0.22 | 0.95 | 74 |
| 1003053 | 5' ETS | C | CG | 0.48 | 0.23 | 0.85 | 51 |
| 1003545 | 5' ETS | C(CCGT)2 | C(CCGT)1 | 0.4 | 0.19 | 0.76 | 71 |
| 1005721 | 5' ETS | G | A | 0.42 | 0.16 | 0.81 | 83 |
| 1006297 | ITS 1 | C | A, CTGA | 0.38 | 0.26 | 0.83 | 60 |
| 1007183 | ITS 2 | (CCGT)1 | (CCGT)2 | 0.42 | 0.19 | 0.93 | 80 |
| 1007812 | ITS 2 | T | C | 0.39 | 0.2 | 0.91 | 72 |
| 1007825 | ITS 2 | C | - | 0.38 | 0.17 | 0.95 | 76 |
| 1008007 | 28S.59 | G | A | 0.39 | 0.17 | 0.99 | 106 |
| 1010121 | 28S.2173 | G | GC | 0.34 | 0.32 | 0.77 | 82 |
| 1010123 | 28S.2175 | G | T, GGT, GCGGGGGT | 0.59 | 0.25 | 0.79 | 84 |
| 1010134 | 28S.2186 | T | - | 0.37 | 0.15 | 0.68 | 54 |
| 1010142 | 28S.2194 | GCC | GC, G | 0.64 | 0.21 | 0.72 | 51 |
| 1010988 | 28S.3040 | A | G | 0.33 | 0.24 | 0.52 | 40 |
| 1010997 | 28S.3049 | C | CCT, CCTCTCT | 0.43 | 0.18 | 0.89 | 95 |
| 1011025 | 28S.3077 | C | T | 0.37 | 0.23 | 0.56 | 59 |
| 1011106 | 28S.3158 | C | G, - | 0.44 | 0.33 | 0.61 | 30 |
| 1013136 | 3' ETS | T | G, TG | 0.51 | 0.27 | 0.8 | 78 |
| 1013422 | IGS | C | T | 0.56 | 0.24 | 0.96 | 75 |

(Prevalent, $pVDF > 0.5$ ; Abundant, $0.33 > iVAF > 0.66$ )

Surprisingly, by organizing the 26 human populations into their five superpopulations, blocks of called variants or even possibly alternative haplotypes emerged (yellow arrows, Figure 2A). These population-uncommon (low pVDF) but intra-individual abundant (higher iVAF) variants (Figure 2C) lends support to the idea that besides intra-individual heterogeneity, there is population-level heterogeneity in rRNA.

Altogether, there is substantial intra- and inter-individual variation in human rDNA, and these exciting findings raise far more questions then they answer.

## Discussion

The idea that ribosomes are homogenous is increasingly being challenged. (16,17) In a land-mark study, knock-out of the murine large sub-unit ribosomal protein (RP) RPL38 resulted in the specific loss of translation of a set of internal ribosomal entry site-dependent HOX genes, but not in global translation.(18,19) In a biochemical analysis of RPs in mouse embryonic stem cells, the stoichiometry of the different RPs varies between isolated monosomes and polysomes.(20) At the transcriptomic level, the relative RP mRNA in yeast varies with growth conditions, and human RP mRNA varies across tissues,(21) and between normal and malignant cells.(22)

These studies have interrogated ribosomal variation at the protein subunit and post-transcriptional modification levels.(23) In contrast, the intra-individual genetic variation, especially variants present across the populations such as 28S.59G>A, offers an enticing new dimension to intra-cellular or intra-individual ribosomal heterogeneity. Since the ’backbone’ of the ribosome is the rRNA, it’s tempting to speculate that differences in the RP-composition and rRNA modifications arise from distinct paralogs of *RNA45S.*

The central role of the ribosome to all translation implies that even small changes to translational efficiency or regulation may have large phenotypic consequences. The paramount biological significance of these genes is underscored by how little we know about their variation in humans or other mammals.

It’s relatively straightforward to include rDNA analysis to existing high-throughput DNA- and RNAseq analysis as demonstrated, but further elucidation of how to meaningfully interpret these variants from a population and evolutionary perspective is necessary.

In particular there is a broad class of human genetic diseases called ribosomeopathies, such as Diamond-Blackfan anemia and X-linked dyskeratosis congenita, caused by mutation to ribosomal protein or ribosomal biogenesis factors (reviewed by Nakhoul et al,(24)). Notably, there are no known genetic diseases caused by mutation or variants in nuclear rRNA, but this may be due to the previous perceived difficulty in studying rRNA genes. Yet classical linkage-analysis of an inherited aminoglycodside-induced deafness pedigree led to the identification of a causative mutation in the the 12S mitochondrial rRNA gene, *MTRNR1*.(25)

Intra- and inter-individual heterogeneity of human ribosomal RNA raises fundamental questions about translational variation at three levels; differences amongst individuals or populations in a species; differences between tissues; and differences between each individual ribosome within cells. Ultimately, this study scratches the surface of this topic but to facilitate rapid and open exploration of the consequences of these findings, my laboratory notebooks, data and scripts/methods are unrestrictedly available online (http://rRNA.ca). I invite interested scientists to cooperate in understanding the full implications of this variation in rRNA and the impact this has on human health.

To date I haven’t found evidence for a human rDNA-specific transposable element such as R1- or R2.

## Materials and Methods

An electronic laboratory notebook is available online (http://rRNA.ca) with line-by-line annotated scripts and methods to replicate these experiments in their entirety.

### Reference Genomes and rDNA repeats

The 43 kb reference rDNA repeat, U13369.1 was downloaded from NCBI. The first nucleotide of this sequence is the transcription start site (+1) of the 13.5 kb 45S pre-rRNA, and the last 1 kb is the rDNA upstream promoter region. Initial analyses were hindered by the sequence complexity of the intergenic spacer, so this study focuses on the genic portion of rDNA. PCR-free whole genome sequencing data from CEPH-1463 was aligned to *RNA45S* and its promoter as well as a single copy of the 5S locus from hg38 (chr1:228744112-228746352). The sequences were manually edited to match the consensus supported by the data, with the reasoning that this represents the common rDNA sequence at each position. Three difficult to align regions were masked with ’N’ nucleotides, an AluY element and low complexity sequence in the 5S locus and a simple-repeat sequence in the ITS1. The resulting sequence, *hgr1,* was used in subsequent analysis {Sequence Accession Pending}.

### Datasets

A complete list of sequencing libraries used in this study is available in Supplementary Table 1. Four genomes from each of the 26 human populations were chosen randomly from the 1000 genomes project data(15). I required paired-end Illumina sequencing with a minimum of 20 million sequenced fragments per library. On average the libraries used had 67.8 million sequenced fragments.

The evolutionary Shannon information for each mature rRNA base is based on a mutliple-sequence alignment of rRNA sequences from all domains of life(26) and downloaded from the RiboVision suite.(27)

### Alignment

Initial alignment parameters were selected to maximize sensitivity. DNAseq alignment was performed with *bowtie2 (28)* using the command ‵*bowtie2 --very-sensitive-local -x hgr1 -1 <read1.fq.gz> -2 <read2.fq.gz>*‵. The majority of the alignments were ran on Amazon EC2 using an C4.2xlarge instance. A publicly available instance image with the necessary software and reference genomes is available: *’crown-161229’*.

### Variant Discovery

The purpose of this study was to define the major, high frequency variants in rDNA. Pseudogenized rDNA is a potential source of error in this study, but I reasoned that pseudo-rDNA mutations at identical positions between copies would be rare relative to total rDNA copy number. For variant-discovery, the 107 *hgr1-* aligned bam files were analyzed with the Genome Analysis Toolkit (v.3.6) *HaplotypeCaller (29)* using standard parameters. *HaplotypeCaller* performs local de-novo assembly over variant positions, which makes it well suited for genotyping rDNA variants. The 926 called variants had an alternative allele quality score of >30 PHRED (p < 0.001) and 849 of those variants had a score of >100 PHRED (p < 10^−10^). The type of substitutions and indel size distribution was plotted with bcftools.(30)

This analysis should be interpreted as a conservative, lower bound on rDNA variation. With a larger data-set and more importantly, deeper and less PCR-bias genome sequencing, multi-ploidy variant analysis should yield a more complete set of variants at lower intra-individual variant allele frequencies.

### Measurements of Variant Allele Frequencies

VCF files were parsed into *R* with *vcfR (31)* and population statistics calculated with custom scripts. For the interrogated variant positions, intraindividual reference allele frequency was calculated by the number of reference genotype read coverage, divided by the total read coverage used in genotyping. The inverse of the reference allele frequency was defined as the intra-individual variant allele frequency (iVAF). I calculated iVAF in this way to account for positions with multiple variant alleles and to measure iVAF in genomes where variant alleles may not have been called by the software, yet variant reads were present. The mean intra-individual variant allele frequency is simply the average value of iVAF for each genome in which a variant was present.

Inter-individual, or population variant-detection frequency (pVDF) is how many individuals in the population have a measurable variant at a particular position (iVAF > 0.02), divided by the total number of individuals in which that position was sequenced. This normalizes for uneven sequencing coverage in the genomes, especially at high-GC regions.

The product of iVAF and pVDF therefore, is variant allele frequency in the classical sense, taking into account the pool of all rDNA alleles. Although, this makes the assumption that rDNA copy number is the same in each individual which may not be true. (2,32) As such I have refrained from terming any of the variation polymorphisms even though a sub-set (Table 1) are most likely polymorphisms.

## Acknowledgements

I’d like to thank my supervisor Dr. Dixie Mager. Thanks to Katharina Rothe, Andrew Chapman, Peter Hurley, Chris Laver and Davide Pellacani for their helpful discussions and critical reviews of the manuscript.

I was supported by a studentship award from the Natural Sciences and Engineering Research Council of Canada. Computing resources were a kind gift from hackseq.

## Supplementary Files

*Supplementary Table 1: Supplementary_Table1.xls.*

Sequencing datasets and their accessions used in this study.

*Supplementary Sequence 1: hgr1.fa.gz.*

*Hgr1* reference rDNA sequence. Deposition into online nucleotide database pending.

*Supplementary File 1:107genomes.hgr1.g.vcf.gz.*

*GVCF file of the 926 variants positions in the 107 genomes included in this study.*

